# Investigate Inhibitory Effects of Ginger polyphenols compare to Simvastatin towards HMG-CoA reductase: An Integrated Molecular Docking and Molecular dynamic simulation

**DOI:** 10.1101/2021.08.11.455913

**Authors:** Aweke Mulu Belachew, Asheber Feyisa, Mulugeta Gajaa Ufgaa, Teslim Yimama Yesuf

## Abstract

Diabetes is an increasing problem in Ethiopia, affecting up to 6.5% of Ethiopian adults. There are serious complications associated with diabetes including macrovascular and microvascular. Controlling Lipid profiles and blood glucose significantly reduces the risk of complications. Statins are the only current treatment for both type 1 diabetes and Type 2 diabetes dyslipidemia. HMG-CoA reductase plays a central role in the production of cholesterol which, associated with cardiovascular disease (CVD). Statins have been found to reduce cardiovascular disease and mortality in those who are at high risk. Nonetheless, it has adverse effect, such as drug-related hypoglycemia and high cost. These situations lead to develop suitable phytotherapeutic agents with less frequent side effects. Ginger (Zingiber officinale) is widely consumed as a spice, and numerous studies suggest that ginger may have beneficial effects for diabetes and dyslipidemia. But, further studies are needed to investigate effects of binding affinity and binding site residues for major ginger extract polyphenols towards target HMG-CoA reductase. In this study, ADMET web server, Auto-Dock 5.4 and Gromacs 2020 were used. Out of eleven major gingers polyphenols screened three selected based on docking energy compare to Simvastatin for MD simulation. The predicted binding affinity for 6-paradol, 6-shogaol and gingerdione were −8.51, −6.93, −9.24 kcal/mol, respectively. The results of molecular dynamic simulation are consistence with docking. The predicted ligand binding site residues are Arg641, Gly808, Arg641, Met781, Ser794 and Arg595. In conclusion, 6-paradol, 6-shogaol and gingerdione could be possible therapy because, of interactions with target HMG-CoA reductase. Therefore, further wet lab study will be needed, for the better understanding of the mechanism of action of ginger extract by which it modulates liver and kidney vivo condition.

## Introduction

Diabetes mellitus is the most common metabolic and endocrine disorder worldwide. It is linked to disturbances in carbohydrate, fat and protein metabolism and important study area because of the global prevalence of diabetes is projected to rise in coming years ^[1]^. It is an increasing problem that contributes significant mortality from increased microvascular and macrovascular complications. Recently, World Health Organization (WHO) reported that 422 million people around the world suffer from diabetes, that this number will double by 2030, and that 80% of diabetics live in developing countries ^[2 6]^. Particularly, the number of diabetic cases in Ethiopia in 2019 was estimated at 19.4 million and by 2045 this figure is expected to rise to 47.1 million. And also, an estimated 700 million adults worldwide will have diabetes by 2045 ^[3 6]^. Over the last 30 years, type 2 diabetes has changed from relatively mild ailment associated with aging to one of a major cause of premature mortality and morbidity in most countries ^[5 6]^. The clinical presentation revealed that diabetes is associated with a cluster of interrelated plasma lipid and lipoprotein abnormalities, including reduced HDL cholesterol, predominance of small dense LDL particles, and elevated triglycerides ^[4]^. Impaired insulin action at the level of the adipocyte is believed to result in defective suppression of intracellular hydrolysis of TGs with the release of free fatty acids into the circulation ^[2]^. Treatment for diabetic hyperlipidemia includes lifestyle interventions, smoking cessation and drugs such as Simvastatin ^[6 7]^. But, lipid-lowering drugs are associated with serious adverse effects ^[8 9]^. Recently, there has been a focus on the search for new drugs capable of dietaries and medicinal plants as dietary components with cholesterol-lowering potential are considered to be useful in preventing dyslipidemia ^[10 11 12]^. So far, several studies have been reported hypoglycemic effects of phytochemicals by using both animal models and human beings ^[8 12 13 14]^. But, still we need lipid-modifying active compounds are required to achieve significant improvement in the lipoprotein profile of diabetic patients. Since, HMG-CoA (3-hydroxy-3-methylglutaryl-coenzyme A) reductase (HMGR) catalyzes the rate limiting step in cholesterol biosynthesis ^[15]^ the study designed to investigate the effect of ginger polyphenols on activity of HMG CoA reductase, and to compare the results with the effect of simvastatin.

For the past several years there have been noticeable improvement in research that focuses on investigating the biological activities of ginger extracts with an emphasis on medicinal function ^[12 13 14]^. These have mainly led to the design and development of numerous studies to determine their antioxidant, antibacterial, anticancer, antihyperglycimia, antidislipidiamia, analgesic, anti-inflammatory, hypoglycemic, other pharmacological activities and antipyretic properties and also received extensive attention as a botanical dietary supplement in the Ethiopia, China, India, United States, Europe and other part of the world ^[16 17 18 19 21 23]^. There are relatively few studies on identifying effect of isolated ginger rhizome major extract on diabetes and hyperlipidemia, but the ones that have been done on crude extract suggest that ginger might be a useful natural treatment for diabetes dyslipidemia. Even though, screening studies have been documented the bioactive importance of several major ginger extract, their target interaction remain poorly characterized in diabetes dyslipidemia. Therefore, this study answers the question which ginger extract have better affinity with target HMG CoA reductase. Ginger derived polyphenols are attractive drug candidate, but they are found in tiny amount and extracting and purifying is not feasible. So far, several studies have been reported major bioactive ginger extract including n-gingerol (n=6, 8, 12), n-shogaol (n=6, 8, 12), α-pinene, camphene, bornyl acetate, 2-undecanone, citronellyl acetate, α-copaene, geranyl acetate 6-methyl-5-hepten-2-one, myrcene, α- and β-phellandrene, limonene, 6-paradol, gingerdiones, 6-gingesulfonicacid, 6-hydroxyshogaol and hexahydrocurcumin ^[8 12 20 22 24 25 26]^. The goal of this project is to screen and identify novel HMG CoA reductase inhibitors out of aforementioned ginger polyphenols and to understand the interacting amino acids residues. And also, it was assessed whether ginger extracts are potential enzymes inhibitors as compare to Simvastatin, using molecular docking and molecular dynamic simulation approach. Inhibitors of this enzyme are useful in managing dyslipidemia and in prevention of macrovascular and microvascular complication. In present days, computational techniques offer convenient approach for analysis of possible inhibitor of target molecules in the area medical science ^[27 28 29 30 31 32 33]^. Here, ADMET, Auto-Dock and molecular dynamics (MD) simulations approaches have been utilized by using HMG-CoA reductase as target. Out of eleven major gingers extract three selected based on docking binding affinity and good interaction with the active site residues. The predicted binding affinity for 6-paradol, 6-shogaol and gingerdione were −8.51, −6.93, −9.24 kcal/mol, respectively. The predicted ligand binding site residues are Arg641, Gly808, Arg641, Met781, Ser794 and Arg595. This study could add knowledge about the effects of major ginger extract on dyslipidemia that might be useful in encouraging possible treatment of diabetics in Ethiopia and the world with ginger. It is hoped that ginger and other plants in this family would open up clinical research, trials using humans and biochemical methodologies and possibly provide new treatment or prevention options for diabetes.

## Materials & Methods

### Materials

Computer system (Hp), with the following specification properties; CPU Dual@2.30 GHz, Intel® Core i10-6100U, 200 Gigabyte RAM was used throughout the present study. The software download and installed include Ubuntu 18.04 LTS, Gromacs 2020, Auto-Dock 5.4 software, Avogadro software, Discovery Studio Visualizer v16.1.0.15350, Swiss ADME online software.

### Ligand preparation

Libraries containing about eleven compounds with small molecular weights were used for this study. Chemical structures of eleven major ginger extracts were retrieved from the Pub-Chem database (https://pubchem.ncbi.nlm.nih.gov/). Avogadro software, version 2.4, 2010 ^[37]^, was used for geometrical refining of chemical structures of selected major ginger extracts by applying GAFF force field with 5000 steps and steepest descent algorithm, which was later saved as Protein Data Bank (PDB) file format. In order to screen drug likeness the SMILES format compounds were filtered through Lipinski’s rule of five and ADMET prediction using AdmetSAR 2.0 webserver ^[38]^ and swissADMET webserver to computed blood brain barrier (BBB) permeant, gastro-intestinal (GI) absorption, cytochrome inhibition. Finally, based on ADME prediction, ligands were made ready for molecular docking analysis using Auto Dock 4.2.

### Protein preparation

Three-dimensional coordinates of HMG-CoA reductase (PDB ID: 1HW9) with RMSD 2.33 Å was obtained in .pdb format from the Protein Data Bank (https://www.rcsb.org/). Structural evaluation and stereo chemical quality of the protein structure was checked through the Ramachandran plot analysis by using UCLA-DOE server-web to assess the overall quality and local quality analysis, based on the Z-score and knowledge-based energy calculations ^[39]^. Validate the modeled structure using the UCLA-DOE server (http://servicesn.mbi.ucla.edu/) ^[18 19 20]^. The modeled 3-D structure was then validated and confirmed by using the RAMPAGE, ERRAT, and Verify 3-D online servers. Based on UCLA-DOE server-web results missed residues and atoms were fixed by using Swiss-pdb viewer and rescreened Structural evaluation and stereo chemical quality ^[39]^. Finally, the .pdb file was entered into Auto Dock 4.2 for preparation of a .pdbqt file and grid box creation. Water molecules and other atoms were excluded, and ADT measured the Gasteiger charges for protein atoms; Auto Grid was used with a grid box to create the grid map. The size of grid was determined at 106 × 126 × 126 xyz points with a grid spacing of 0.5Å and a grid Centre at dimensions (x, y, and z, respectively): −6.81, 6.528, and 0.861was designated.

### Molecular docking

Candidate ginger extracts were individually docked into the binding site of HMG-CoA reductase by using Auto-Dock 5.4. First, polar hydrogen was added to ginger extracts and Simvastatin followed by Gasteiger partial atomic charges were added. Secondly, HMG-CoA reductase was prepared by adding the polar hydrogen, united atom Kollman charges and the protein structure was saved in. pdbqt format that includes atomic partial charges atoms. Last, docking was performed using Lamarckian genetic algorithm with maximum number of evaluation with generation of 27,000 and with the 250 as the population size and with the default values for other parameters ^[40]^. Upon successful molecular docking simulation, the best conformation with lowest binding energy was extracted from the population of 20 individuals. At the end, the top score poses were evaluated by using Ligplus+ software ^[41]^ to visualize 2-D protein-ligand interactions that includes the hydrogen bond and hydrophobic interaction between the ligand and main chain or side chain elements of the protein.

#### Molecular dynamics (MD) simulations

Molecular Dynamics (MD) simulation of the free HMG-CoA reductase and docked HMG-CoA reductase were performed using GROMACS 5.1.2 package for 100 ns at 300 K temperature. Entire simulations are carried out using GAFF force field ^[42]^. The System was placed at center of cubic box with a dimension of 1.2 nm and the systems was solvated with TP3P water model and charges of the system was neutralized upon adding Na^+^ and Cl^−^ with 0.1 M ionic strength. Energy minimization was performed by steepest descent method to the maximum gradient of 1000 kJ/mol/nm to remove bad geometry. Before production MD simulations, the minimized system was heated with constant temperature for 5000ps. Then the system was coupled under constant external stress and at finite temperature, with Parrinello-Rahman barostat to equilibrate at 1 bar pressure for 2000ps. The non-bonded term interactions are calculated using short-range cut-off and the long-range cut-off for columbic and Lenard-Jones interactions and bonds associated with hydrogen atoms were constrained based on the LINCS protocol ^[43]^. After successful completion of system equilibration, production MD run was carried out for 100 ns and trajectory structures were stored at every 10ps. The trajectory files were visualized by using VMD ^[44]^.

## Results

## Discussion

Lipids metabolism is usually raised and play an important role in the pathogenesis of Macro vascular and Micro vascular in the diabetes mellitus patients ^[15]^. The current study revealed that the crude ginger extract decrease Lipids profiles both in animal model and Human ^[16 17 18 19]^. Even though, subjective side effects due to ginger powder were significantly less than Simvastatin no study address this issue in silico study. Research findings have shown HMG-CoA reductase highlighted as simvastatin target HMG-CoA reductase in dyslipidemia during Diabetes ^[15]^. But, few study reported a comparison of simvastatin efficacy with ginger rhizome extract to show similarity and difference amongst the two to manage dyslipidemia. At the present time, several studies have been revealed the significant out of using computational approaches to investigate small molecules-target interactions ^[29 30 31 32 33]^. In this study, we applied molecular docking, molecular dynamics (MD) simulations, and other computer aided drug design (CADD) methods to study HMG-CoA reductase and ginger extract. The aim of this study was to assess whether ginger extracts are more potential HMG-CoA reductase inhibitors than simvastatin, using a molecular docking and molecular dynamic simulation by using study. At the beginning of study, top eleven selected Ginger rhizomes extract and Simvastatin were subjected to ADMET properties and Lipinski’s rule 5 prediction and the results are tabulated (Table 1 and Table 2). The pharmacokinetic properties and Lipinski’s rule 5 of top-three compounds close to co-crystal reference drug as shown in Table 1 and Table 2. In this study, reference drug consist of 7 torsional angles in its structure, while the three ginger extract consist of rotatable bonds in the range of 9-14. The Hydrogen bond acceptor (HBA) and donor (HBD) profiles of three selected ginger extract close to reference drug. ADMET for top-five compounds (Hexahydrocurcumin, 10-shogaol, 10-gingerol, 8-shogaol and 6-shogaol) close to reference drug as shown in Table 2. Among screened compounds, three passed the ADMET prediction ESOL Solubility (mg/ml, GI absorption, BBB, P-gp substrate, a CYP1A2 inhibitor, CYP2C19 inhibitor, CYP2C9 inhibitor, CYP2D6 inhibitor and CYP3A4 inhibitor with high GI absorption and high BBB permeability. Simultaneously, before going for docking, three-dimensional crystal structure of the HMG-CoA (3-hydroxy-3-methylglutaryl-coenzyme A) reductase (PDB Id: 1hw9) was analyzed by using PRO-CHECK web server ^[15]^. The Ramachandran plot statistics demonstrated the stereo chemical quality of HMG-CoA reductase (PDB Id: 1hw9) is in good quality; 90.0 % of amino residues located within the most favored and additional allowed regions and only 0.1% of the amino acids found in the disallowed region (**Figure. S2**), which is in acceptable limit. The calculated overall average G-factors score of dihedral angles and main chain covalent forces for the HMG-CoA reductase was found to be 0.33. Moreover, the calculated PROSA Z-score value of 2.24 shows that HMG-CoA reductase is found to be within the range of native conformations of experimental structures of similar size, thus it is good quality and reliable. Significant negative energies were observed in the local HMG-CoA reductase quality plot suggesting that it possess unproblematic and un-erroneous.

**Table 1.** Molecular pharmacokinetic properties of co-crystal Simvastatin and top-five ginger extract compounds computed using admetSAR 2.0 webserver

| Molecule | [6]- paradol | 6-shogaol | Gingerdione | Hexahydrocurcumin | Simvastatin |
| --- | --- | --- | --- | --- | --- |
| MW(g/mol) | 278.39 | 348.55 | 292.37 | 374.43 | 418.57 |
| Fraction Csp3 | 0.59 | 0.55 | 0.53 | 0.38 | 0.76 |
| Rotatable bonds | 10 | 11 | 10 | 10 | 7 |
| HBA | 3 | 3 | 4 | 6 | 5 |
| HBD | 1 | 0 | 1 | 3 | 1 |
| TPSA (Å<sup>2</sup>) | 46.53 | 35.53 | 63.60 | 96.22 | 72.83 |
| XLOGP3 | 4.11 | 6.08 | 2.88 | 2.70 | 4.68 |
| ESOL Log S | 3.65 | -6.35 | -3.02 | -3.53 | -4.92 |

**Table 2.** ADMET properties of co-crystal Simvastatin and top-five ginger extract compounds computed using AdmetSAR webserver

| Molecule | [6]- paradol | 6-shogaol | Gingerdione | Hexahydrocurcu min | Simvastatin |
| --- | --- | --- | --- | --- | --- |
| Log S solubility (mg/ml) | -3.72 | -6.61 | -3.88 | -4.37 | -5.94 |
| GI absorption | High | High | High | High | High |
| BBB permeant | Yes | Yes | Yes | No | No |
| P-gp substrate | No | No | No | Yes | No |
| CYP1A2 inhibitor | yes | Yes | Yes | No | No |
| CYP2C19 inhibitor | No | No | No | No | No |
| CYP2C9 inhibitor | No | No | No | No | Yes |
| CYP2D6 inhibitor | yes | Yes | Yes | Yes | No |
| CYP3A4 inhibitor | No | Yes | No | Yes | Yes |
| Log Kp (cm/s) (skin permeation) | -5.08 | -4.11 | -6.04 | -6.67 | -5.53 |

Next, we studied the affinity of three ginger rhizome extract and simvastatin which have strong drug-like properties. As shown in Table 3 the affinity (Kcal/mol), hydrogen bonds, and binding residues of top three ginger rhizome extract and simvastatin. We found that three ginger rhizome extracts have strong binding affinity with HMG-CoA reductase and inhibitor constant compare to simvastatin. The three ginger extract includes, 6-paradol −6.93 (9.88pM), 6-Gingerdione −8.51 (575.41pM) and 6-shogaol −9.24 (169.42pM) kcal/mol, but the Ki values are significantly larger, compared to simvastatin (Table 3). In comparison with all three ginger extract and simvastatin, 6-shogaol had the highest binding energy and 6-Gingerdione has the least Ki values indicating that Simvastatin can capable of inhibiting the activity of human HMG-CoA reductase protein even at 575.41pM concentration and rest of the derivatives of ginger extract had more Ki values in the range of 225.91–780.00pM concentration (Table 3 supportive). The molecular interaction revealed 6-Gingerdione, 6-shogaol and 6-gingerol to possess inhibitory potential against enzymes with binding energy range of −□8.84 to −9.24 kcal/mol, and Ki values of 169.42-575.41μM (Table 3). Our study suggests that 6-Gingerdione, 6-shogaol and 6-gingerol satisfies in silico parameters tested and is expected to efficiently inhibit cholesterol *biosynthesis* pathway more than simvastatin. Furthermore, as shown in table 3 the number of hydrogen bonds between statins and amino acid residues of HMG-CoA reductase revealed in the range of 3-5 for ginger extract and 7 for simvastatin, while other ginger extract had less than three hydrogen bonds (**Table S3 and S1**). These results indicate, based upon the binding energy of simvastatin, and three ginger extract, that statins could be efficient HMG-CoA reductase inhibitors. The Ligplot+ results for all the eight ligands are shown in figure 1. The predicted ligand binding site residues comprises Arg641, Gly808, Arg641, Met781, Ser794 and Arg595 as favorable binding site region to dock the ligands (figure 1). Except 6-paradol, all other ginger extract interacting with the residues Arg641, Arg641, Met781, Ser794 and Arg595, which indicates the importance of these residues to maintain HMG-CoA reductase-ginger extracts complex (Table 3). Except, all the ginger extracts capable of forming hydrogen bonds with HMG-CoA reductase (Table 3 supportive). Overall molecular docking study showed that the compound simvastatin had highest binding affinity that can produce the inhibition even at low concentration, hence, we have prioritized simvastatin as the best candidate for further optimization for in-vitro studies.

**Table 3.** The calculated binding energy, hydrogen bonds and contacting HMG-CoA reductase residues of co-crystal Simvastatin and top-three ginger extract compounds using Auto-Dock 4.5 structure

| Name | Binding energy<br>[kcal/ mol] | Ki uM (micro<br>molar) | RMSD<br>(A<sup>0</sup>) | Hydrogen<br>Bonds | Contacting Enzymes<br>Residues |
| --- | --- | --- | --- | --- | --- |
| Simvastatin | -6.92 | 8.43 | 22.104 | 8 | Glu559, Lys736, Asn755,<br>Arg590, Ser684, Asp690,<br>Ser684 and Lys691 |
| 6- paradol | -6.93 | 9.88 | 42.507 | 1 | Gly808 |
| 6-Gingerdione | -8.51 | 575.41 | 21.335 | 2 | Arg641 and Arg595 |
| 6-shogaol | -9.24 | 169.42 | 35.658 | 3 | Arg641, Met781, Ser794 |

**Figure 1.**
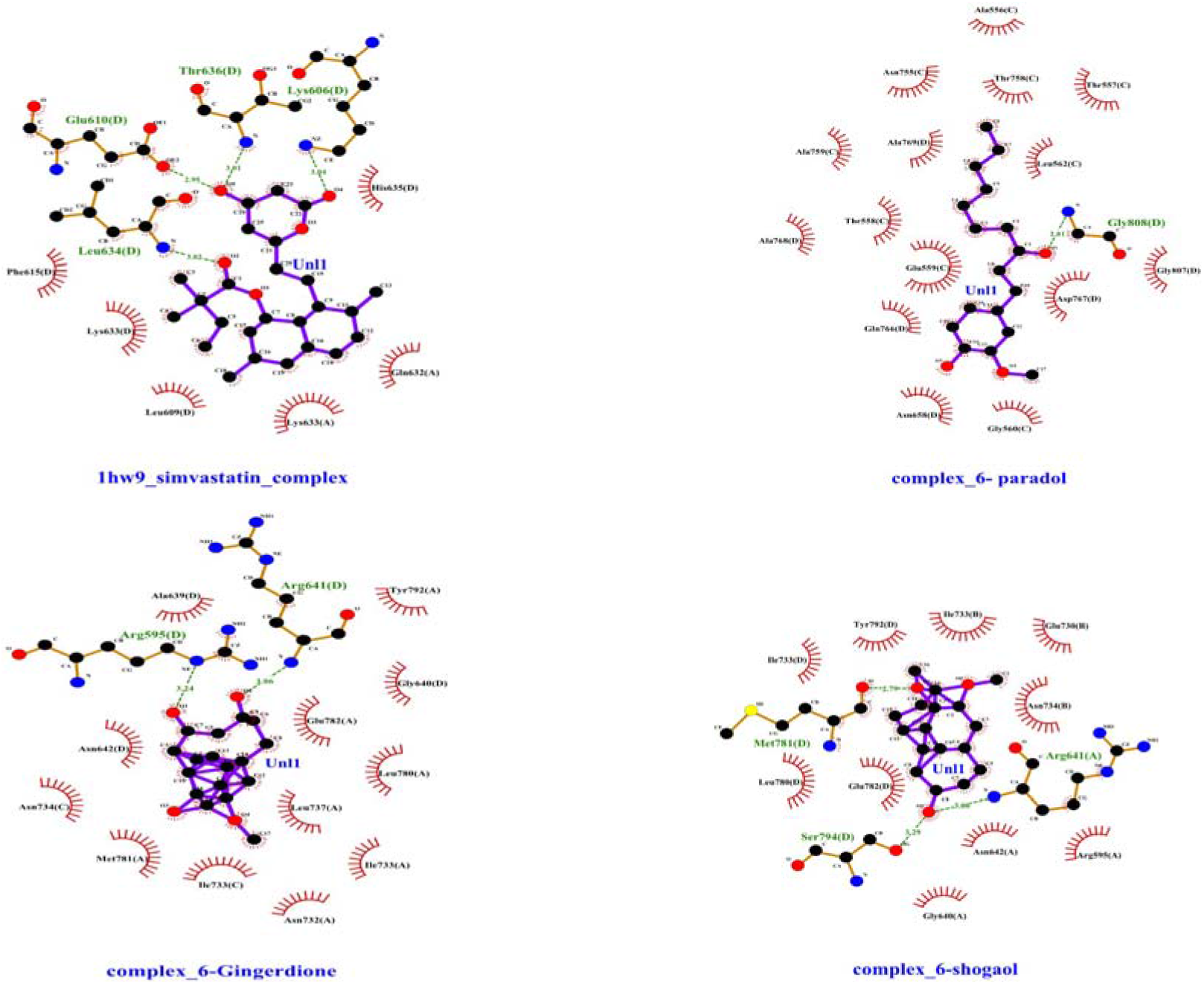
Ligplot+ diagram for the docked Simvastatin, 6-paradol, 6-Gingerdione and 6-shogaol in the HMG-CoA reductase binding site, showing binding environment and interaction profile. The dashed green lines indicate hydrogen bonds, and the half-moon indicates van der Waals interactions

Based on Auto-Docking result, Molecular dynamics simulations were performed for 100□ns for three HMG-CoA reductase-ginger extracts complex to assess the conformational stability and fluctuations. Here we compare the results of the simvastatin-complex with that of ginger extracts complex. As shown in figure 2 a root mean square deviation (RMSD) graph was obtained by MD simulation for the Cα backbone of complex, and it showed that the complex was stable. These results go a line with previous reports in this journal, showing time-dependences of the MD trajectories was examined using the RMSD, RMSF and Rg of all backbone atoms ^[34 29 30]^. Analysis of Root Mean Square Deviation (RMSD) showed HMG-CoA reductase-gingerdione complex is stable after 20ns simulation while HMG-CoA reductase-6-shogaol complex showed stable conformation after 40ns and then remained stable over 100ns as displayed in Figure 2. However, HMG-CoA reductase-simvastatin and HMG-CoA reductase-paradol showed stable conformation throughout simulation as displayed in Figure 2. The Root Mean Square Fluctuation (RMSF) indicates occurrence of local changes along with the protein chain residues at specific temperature and pressure. Root mean square fluctuation (RMSF) values for all residues in the HMG-CoA reductase-ginger extracts complex were studied as shown in figure 3. The root mean square fluctuation (RMSF) for all atoms of the HMG-CoA reductase-ginger extracts complex shows fluctuations within 0.1–1.2nm. In line with previous studies RMSF is an important parameter that yields data about the structural adaptability of Cα atoms of every residue in the system ^[29 30 31 35]^. RMSF shows the fluctuation of terminal amino acids between 250-446aa, which is consistence with previous study ^[15]^. Last but not least, the radius of gyration (Rg) value of the complex backbone was determined for a 100ns trajectory as shown figure 4, and it was found that the HMG-CoA reductase-ginger extracts complex were stable and densely packed until 40ns. These basic findings are consistent with research showing that, the Rg is used to assess the overall dimensions and stabilities of the enzyme-ligand complex and is a function of the mass-weighted RMS distances of atoms from the center of mass ^[36]^.

**Figure 2.**
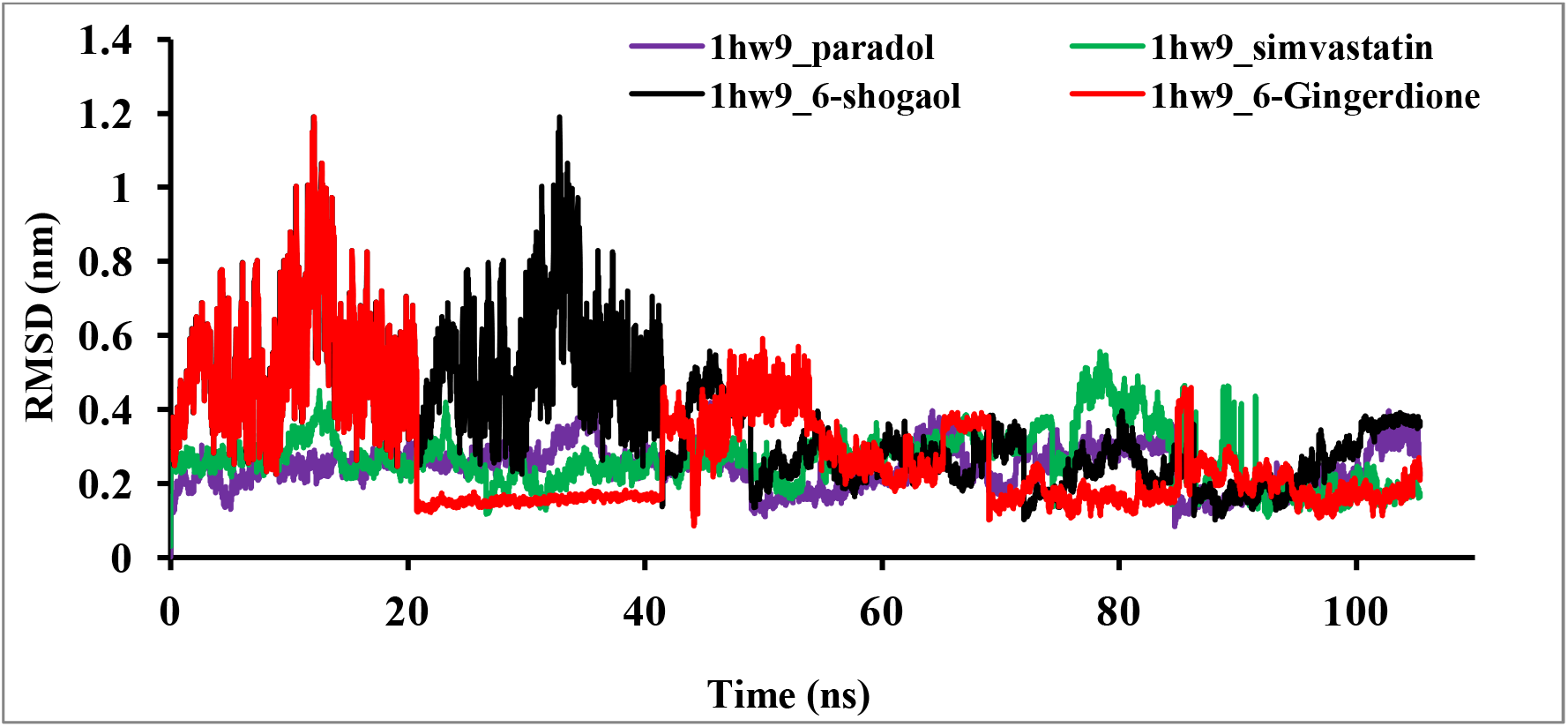
shows conformation changes of HMG-CoA reductase-ligands complex and the root mean square deviation is calculated for 100 ns at 300 K simulation.

**Figure 3.**
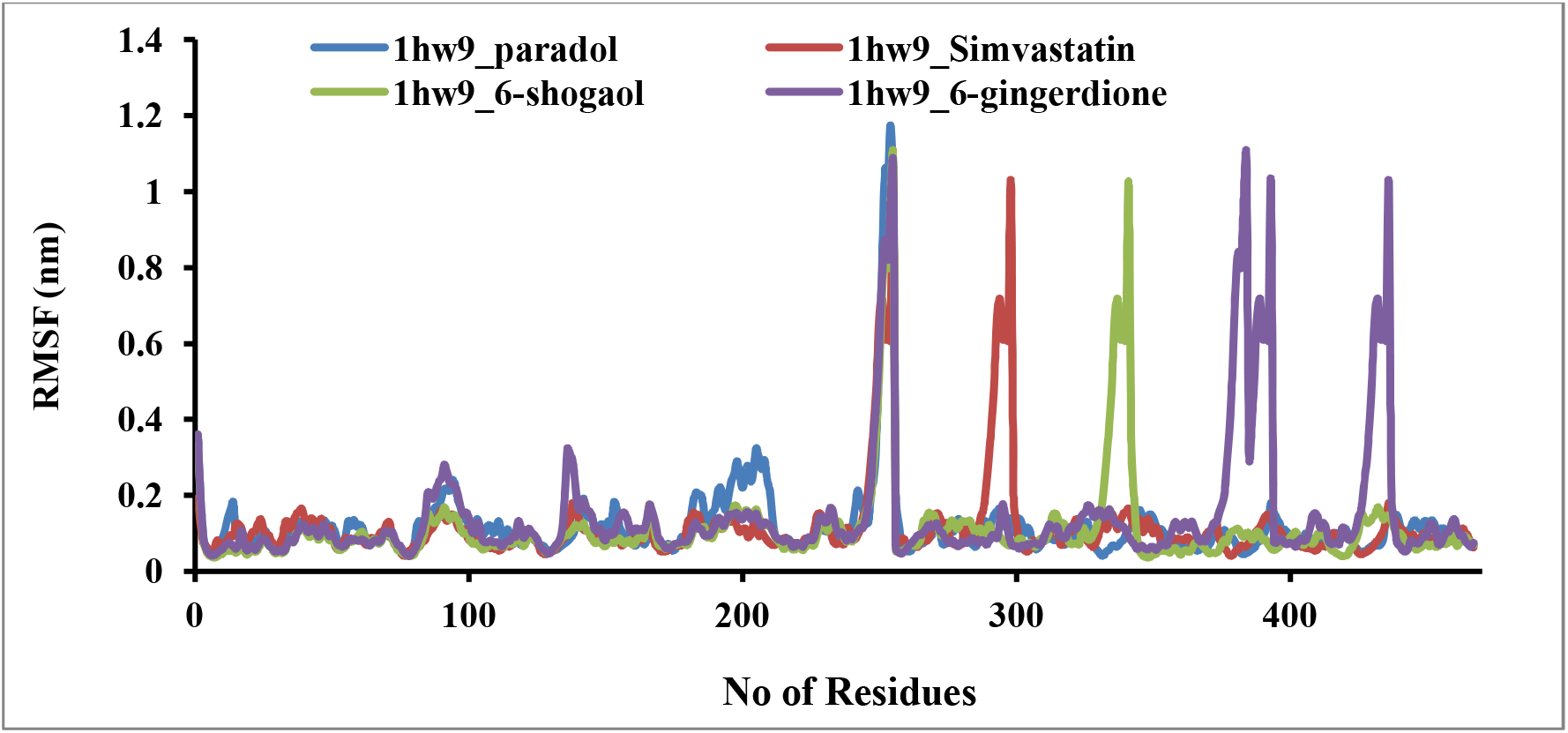
shows conformation changes of HMG-CoA reductase-ligands complex and the root mean square fluctuation is calculated for 100 ns at 300 K simulation.

**Figure 4.**
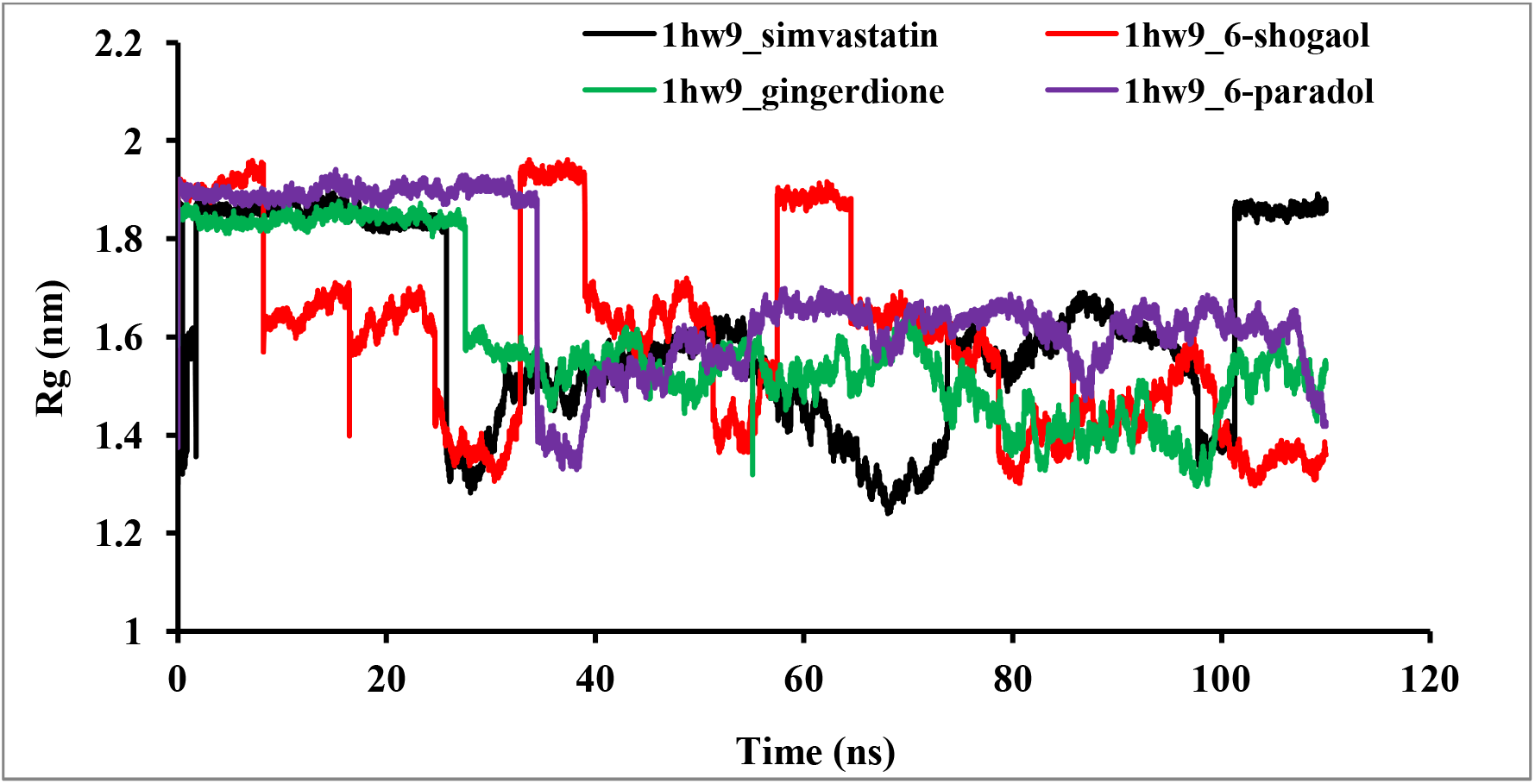
shows conformation changes of HMG-CoA reductase-ligands complex and the radius of gyrations are calculated for 100 ns at 300 K simulation.

### Conclusions

From these research reports one might predict that lowering the lipids profile would prevent the progression of plague formation. However, to date, lipids profile-specific drug is not available and the effect of routinely used cardiovascular drugs on HMG-CoA reductase is not known. Hence, it is very essential to study the association between known cardio-protective drugs such as simvastatin and HMG-CoA reductase. To our knowledge, this is the first study which has proposed the three-dimensional structure of HMG-CoA reductase with possible binding site residues and predicted the binding efficiency of ginger extracts towards HMG-CoA reductase. The proposed HMG-CoA reductase can serve as a good starting point to investigate the interactions with ginger extract as well as simvastatin. Consequently, ginger extracts are a favorable choice for treatment of dyslipidemia compared with simvastatin. Therefore, it is suggested a more extensive wet lab study which can measure the effectiveness of various doses of ginger-based medications with differing types and severities of dyslipidemia is examined. Furthermore, the average RMSD values of the backbone atoms in docked ginger extracts were calculated from 100ns and showed stable RMSD values between after 20ns and 40ns for HMG-CoA reductase-gingerdione complex and HMG-CoA reductase-6-shogaol complex, respectively at reasonably consistent temperature (∼300 K) and pressure (1bar), whereas HMG-CoA reductase-simvastatin and HMG-CoA reductase-paradol complex showed stable RMSD values through tout simulation with same cut-off parameters. To completely investigate the effects of selected ginger extracts, this study recommended the roles of extracts identified for further exploration in a wet lab experiment.

## Acknowledgement

For the success of this work we would like to acknowledge effort of our family for their inspiration. We acknowledge the university of Addis Ababa Science and Technology for using computational resources. Last but not least I would like acknowledge Ministry of innovation and technology of Ethiopia for providing computation power for performing MD simulations.

## References

1. Rains J.L and Jain S.K. (2011). Oxidative stress, insulin signaling, and diabetes. Free. Radic. Biol. Med; 50:567–575. Doi: 10.1016/j.freeradbiomed.2010.12.006.

2. Gupta R, Ghosh A, Singh A.K and Misra A. (2020). Clinical considerations for patients with diabetes in times of COVID-19 epidemic. Diabetes Metab Syndr; 14(3):211–212.

3. Cho NH, Shaw JE, Karuranga S, Huang Y, da Rocha Fernandes JD, Ohlrogge AW and Malanda B. (2020). IDF Diabetes Atlas: Global estimates of diabetes prevalence for 2017 and projections for 2045. Diabetes Res Clin Pract; 138:271–281.

4. American Diabetes Association. (2020). Classification and Diagnosis of Diabetes: Standards of Medical Care in Diabetes—2020. Diabetes Care 2020 Jan; 43(1): S14–S31

5. Singh GM, Danaei G, Farzadfar F, Stevens GA, Woodward M, Wormser DK, et al. (2013). The age-specific quantitative effects of metabolic risk factors on cardiovascular diseases and diabetes: a pooled analysis. Plos One; 8 :(7) e65174.

6. Saeedi P, Petersohn I, Salpea P, Malanda B, Karuranga S, Unwin N, Colagiuri S, Guariguata L, A Motala A et al. (2019). Global and regional diabetes prevalence estimates for 2019 and projections for 2030 and 2045: results from the international diabetes federation diabetes atlas, 9th edition. Diabetes Research and Clinical Practice; 157:107843.

7. Collins AJ, N Foley R, T Gilbertson D and Chen SC. (2015). 2014 USRDS annual data report: Epidemiology of kidney disease in the United States. United States Renal Data System. National Institutes of Health, National Institute of Diabetes and Digestive and Kidney Diseases. Kidney Int Suppl; 5(1): 2–7.

8. Singh AB, Singh NA, Maurya R and Srivastava AK. (2009). Anti-hyperglycaemic, lipid lowering and anti-oxidant properties of [6]-gingerol in db/db mice. Int. J. Med. Med. Sci;1:536–544

9. Spiller HA and Sawyer TS. (2006). Toxicology of oral antidiabetic medications. Am. J. Health Pharm; 63:929–938.

10. Gao Y, Zhang RR, Li JH, Ren M, Ren ZX, Shi JH, Pan QZ and Ren SP. (2012). Radix Astragali lowers kidney oxidative stress in diabetic rats treated with insulin. Endocrinology; 42:592–598.

11. Klein G, Kim J, Himmeldirk K, Cao Y and Chen X. (2007). Antidiabetes and Anti-Obesity Activity of Lagerstroemia speciosa. Evidence Based Complement. Altern. Med; 4:401–407.

12. Kim CY, Seo Y, Lee C, Park GH and Jang JH. (2018). Neuroprotective effect and molecular mechanism of [6]-gingerol against scopola-mine-induced amnesia in C57BL/6 mice. Evidence Based Complement. Altern. Med; 2018:8941564.

13. Yagihashi S, Miura Y and Yagasaki K. (2008). Inhibitory effect of gingerol on the proliferation and invasion of hepatoma cells in culture. Cytotechnology; 57:129–136.

14. Chakraborty D, Mukherjee A, Sikdar S, Paul A, Ghosh S and Khuda-Bukhsh AR. (2012). [6]-Gingerol isolated from ginger attenuates sodium arsenate induced oxidative stress and plays a corrective role in improving insulin signaling in mice. Toxicol. Lett; 210:34–43.

15. Istvan SE and Deisenhofer J. (2002). Structural mechanism for statin inhibition of 3-hydroxy-3-methylglutaryl coenzyme A reductase. Am Heart J; 144(6): S27–32.

16. Tung BT, Thu DK, Thu NTK and Hai NT. (2017). Antioxidant and acetyl cholinesterase inhibitory activities of ginger root (Zingiber officinale Roscoe) extract. Journal of Complementary and Integrative Medicine; 14(4):1–10.

17. Shidfar F, Rajab A, Rahideh T, Khandouzi N, Hosseini S and Shidfar S. (2015). The effect of ginger (Zingiber officinale) on glycemic markers in patients with type 2 diabetes. Journal of Complementary and Integrative Medicine; 12(2):165–170.

18. Ezzat SM, Ezzat MI, Okba MM, Menze ET and Abdel-Naim AB. (2018). The hidden mechanism beyond ginger (Zingiber officinale Rosc.) potent in vivo and in vitro anti-inflammatory activity. Journal of Ethno pharmacology; 214:113–123.

19. Ansari M, Porouhan P, Mohammadianpanah M, Omidvari S, Mosalaei A, Ahmadloo N, Nasrollahi H and Hasan Hamedi S. (2016). Efficacy of ginger in control of chemotherapy induced nausea and vomiting in breast cancer patients receiving doxorubicin-based chemotherapy. Asian Pacific Journal of Cancer Prevention; 17(8):3877–3880.

20. Park M, Bae J and Lee DS. (2008). Antibacterial activity of [10]-gingerol and [12]-gingerol isolated from ginger rhizome against periodontal bacteria. Phytotherapy Research; 22(11):1446–1449.

21. Chakotiya AS, Tanwar A, Narula A and Sharma RK. (2017). Zingiber officinale: its antibacterial activity on Pseudomonas aeruginosa and mode of action evaluated by flow cytometry. Microbial Pathogenesis; 107:254–260.

22. Saha A, Blando J, Silver E, Beltran L, Sessler J and DiGiovanni J. (2014). 6-Shogaol from dried ginger inhibits growth of prostate cancer cells both in vitro and in vivo through inhibition of STAT3 and NF-κB signaling. Cancer Prevention Research; 7(6):627–638.

23. Pashaei-Asl R, Pashaei-Asl F, Mostafa Gharabaghi P, Khodadadi K, Ebrahimi M, Ebrahimie E and Pashaiasl M. (2017). The inhibitory effect of ginger extract on ovarian cancer cell line; application of systems biology. Advanced Pharmaceutical Bulletin; 7(2):241–249.

24. Suk S, Kwon GT, Lee E, Jang WJ, Yang H, Kim JH, Thimmegowda NR, Chung M, Kwon JY, Yang S, K Kim J, Yoon Park JH and Won Lee K. (2017). Gingerenone A, a polyphenol present in ginger, suppresses obesity and adipose tissue inflammation in high-fat diet-fed mice. Mol. Nutr. Food Res; 61:1700139.

25. Wei C, Tsai Y, Korinek M, Hung P, El-Shazly M, Cheng Y, Wu Y, Hsieh T and Chang F. (2017). 6-Paradol and 6-shogaol, the pungent compounds of ginger, promote glucose utilization in adipocytes and myotubes, and 6-paradol reduces blood glucose in high-fat diet-fed mice. Int. J. Mol. Sci; 18:168.

26. Bernard MM, McConnery JR and Hoskin DW. (2017). [10]-Gingerol, a major phenolic constituent of ginger root, induces cell cycle arrest and apoptosis in triple-negative breast cancer cells. Exp. Mol. Pathol; 102:370–376.

27. Cheng F, Li W, Zhou Y, Shen J, Wu Z, Liu G, W. Lee P and Yun Tang. (2012). AdmetSAR: a comprehensive source and free tool for evaluating chemical ADMET properties. J. Chem. Inf. Model; 52(11): 3099–3105.

28. Tao Y, Li W, Liang W and Van Breemen RB. (2009). Identification and quantification of gingerols and related compounds in ginger dietary supplements using high-performance liquid chromatography-tandem mass spectrometry. J Agric Food Chem; 57(21):10014–21.

29. Bandaru S, Alvala M, Nayarisseri A, Sharda S, Goud H, Mundluru HP and Kumar Singh S. (2017). Molecular dynamic simulations reveal suboptimal binding of salbutamol in T164I variant of β2 adrenergic receptor. PLoS ONE; 12(10): e0186666.

30. Aghili Z, Taheri S, Zeinabad HA, Pishkar L, Saboury AA, Rahimi A and Falahati M. (2016). Investigating the Interaction of Fe Nanoparticles with Lysozyme by Biophysical and Molecular Docking Studies. PLoS ONE; 11(10): e0164878.

31. Ashfaq UA, Saleem S, Masoud MS, Ahmad M, Nahid N, Bhatti R, Almatroudi A and Khurshid M. (2021). Rational design of multi epitope-based subunit vaccine by exploring MERS-COV proteome: Reverse vaccinology and molecular docking approach. PLoS ONE 16(2): e0245072.

32. Antony P and Vijayan R (2015) Identification of Novel Aldose Reductase Inhibitors from Spices: A Molecular Docking and Simulation Study. PLoS ONE; 10(9): e0138186.

33. Raj S, Sasidharan S, Dubey VK and Saudagar P. (2019). Identification of lead molecules against potential drug target protein MAPK4 from L. donovani: An in-silico approach using docking, molecular dynamics and binding free energy calculation. PLoS ONE; 14(8): e0221331.

34. Sharma N, Sharma M, Shakeel E, Jamal QMS Kamal MA Sayeed U, Khan MKA, Siddiqui MH, Arif JM and Akhtar S. (2018). Molecular interaction and computational analytical studies of pinocembrin for its antiangiogenic potential targeting VEGFR-2: A persuader of metastasis. Med. Chem; 14: 626–640.

35. Haq FU, Abro A, Raza S, Liedl KR and Azam SS. (2017). Molecular dynamics simulation studies of novel beta-lactamase inhibitor. J. Mol. Graph. Model; 74: 143–152.

36. Baig MH, Sudhakar DR, Kalaiarasan P, Subbarao N, Wadhawa G, Lohani M, Khan MK, Khan AU. (2014). Insight into the effect of inhibitor resistant S130G mutant on physico-chemical properties of SHV type beta-lactamase: A molecular dynamics study. PLoS ONE; 9, e112456.

37. Hanwell MD, Curtis DE, Lonie DC, Vandermeersch T, Zurek E & Hutchison GR. (2012). Avogadro: an advanced semantic chemical editor, visualization, and analysis platform. Journal of Cheminformatics; 4: 17 (2012)

38. Yang H, Lou C, Sun L, Li J, Cai Y, Wang Z, Li W, Liu G and Tang Y. (2019). AdmetSAR 2.0: web-service for prediction and optimization of chemical ADMET properties. Bioinformatics; 35(6):1067–1069

39. M. Morris G, S. Goodsell D, S. Halliday R, Huey R, E. Hart W, K. Belew R and J. Olson A. (1998). Automated docking using a Lamarckian genetic algorithm and an empirical binding free energy function. J Comput Chem; 19: 1639–1662.

40. Wallace AC, Laskowski RA and Thornton JM (1996). LIGPLOT: a program to generate schematic diagrams of protein-ligand interactions. Protein Eng; 8, 127–134.

41. Wang J, Wolf RM, Caldwell JW, Kollman PA, Case DA. (2004). Development and testing of a general AMBER force field. Journal of Computational Chemistry; 1157–1174.

42. Cotter G, Kaluski E, Milo O, Blatt A, Salah A, Hendler A, Krakover R, Golick A and Vered Z. (2003). LINCS: L-NAME (a NO synthase inhibitor) In the treatment of refractory Cardiogenic Shock: A prospective randomized study. European Heart Journal; 24(14):1287–1295.

43. Humphrey W, Dalke A and Schulten K. (1996). VMD: Visual molecular dynamics. Journal of Molecular Graphics; 14(1):33–38

