## Supplementary material for "Investigate Inhibitory Effects of Ginger polyphenols compare to Simvastatin towards HMG-CoA reductase: An Integrated Molecular Docking and Molecular dynamic simulation": Computational_Biology-supplemantal.docx

Investigate potential of Simvastatin and Ginger extract against HMG-CoA reductase to control dyslipidemia: an Integrated Molecular Docking and Molecular dynamic simulation

Aweke Mulu Belachew^1*^, [Ashebe](https://peerj.com/user/143408/)r Feyisa^2^, Mulugeta Gajaa Ufgaa^3^, and Teslim Yimama Yesuf ^4^

^1^ College of Applied Science, Addis Ababa Science and Technology University, Addis Ababa, Ethiopia

^4, 5^ College of Biological and Chemical Engineering , Addis Ababa Science and Technology University, Addis Ababa, Ethiopia

^2, 3^ College of Natural and Social science, Addis Ababa Science and Technology University, Addis Ababa, Ethiopia

Corresponding Author:

Aweke Mulu Belachew^1*^

Addis Ababa, Ethiopia, 16417, Ethiopia

***Supplemental files***

**Table S1** Molecular pharmacokinetic properties of screened ginger extract by using admetSAR 2.0 webserver

| Molecule | 6-gingerol | 6- hydroxyshogaol | 6-Gingesulfonic acid | 8-shogaol | 10-gingerol | 10-shogaol | Hexahydrocurcumin |
| --- | --- | --- | --- | --- | --- | --- | --- |
| MW(g/mol) | 294.39 | 292.37 | 358.45 | 304.42 | 350.49 | 332.48 | 374.43 |
| Fraction Csp3 |  | 0.47 | 0.59 | 0.53 | 0.67 | 0.57 | 0.38 |
| Rotatable bonds | 10 | 9 | 11 | 11 | 14 | 13 | 10 |
| HBA | 4 | 4 | 6 | 3 | 4 | 3 | 6 |
| HBD | 2 | 2 | 2 | 1 | 2 | 1 | 3 |
| TPSA (Å^2^) | 66.76 Å² | 66.76 Å² | 109.28 Å² | 46.53 Å² | 66.76 Å² | 46.53 | 96.22 |
| XLOGP3 | 3.23 | 2.51 | 2.67 | 5.08 | 5.27 | 6.16 | 2.70 |
| ESOL Log S | -3.234 | 3.14 |  | -4.40 | -4.59 | -5.11 | -3.53 |

**Table S2** The calculated binding energy, hydrogen bonds and contacting HMG-CoA reductase residues of screened ginger extract compounds using Auto-Dock 4.5 structure

| Name | Binding energy  [kcal/ mol] | Ki uM (micro molar) | RMSD (A^0^) | Hydrogen Bonds | Contacting Enzymes Residues |
| --- | --- | --- | --- | --- | --- |
| 6- hydroxyshogaol | -5.40 | 109.98 | 16.074 | 3 | Val801,Gly560 and Glu559 |
| 8-shogaol | -6.85 | 9.51 | 43.167 | 0 | 0 |
| 10-shogaol | -5.50 | 93.21 | 14.631 | 3 | Ala525, Gly560,Gly656 |
| 6_gingerol | -4.91 | 35.879 | 252.49 |  |  |
| 8-gingerol | -4.83 | 24.159 | 285.99 |  |  |
| 10-gingerol | -4.82 | 22.397 | 129.42 |  |  |
| Hexahydrocurcumin |  |  |  |  |  |

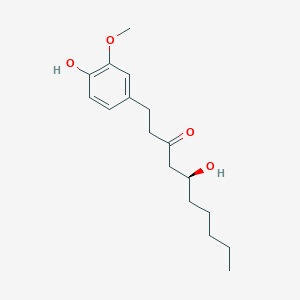

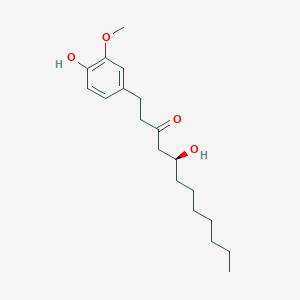

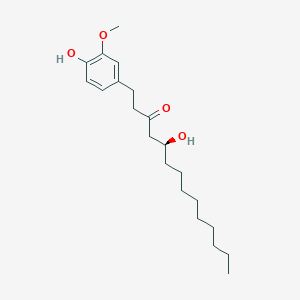

6-gingerol	8-gingerol	10-gingerol

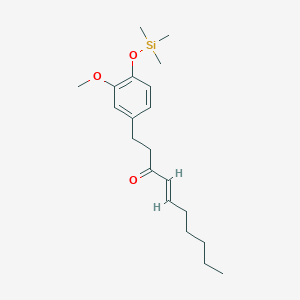

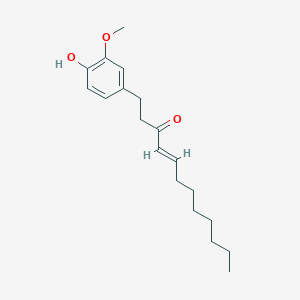

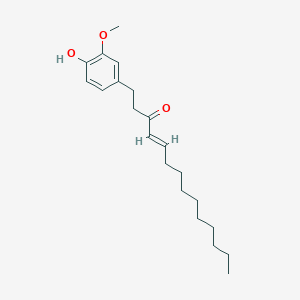

6-Shogoal	8-shogoal	10-shogaol

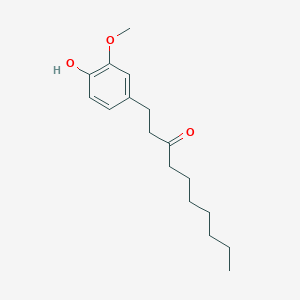

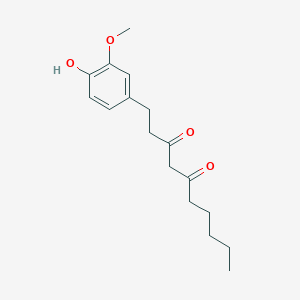

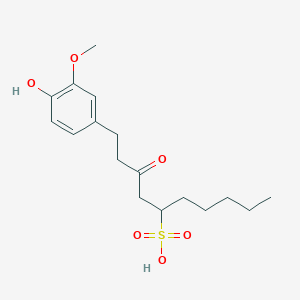

[6]- Paradol	Gingerdione	6-Gingesulfonic acid

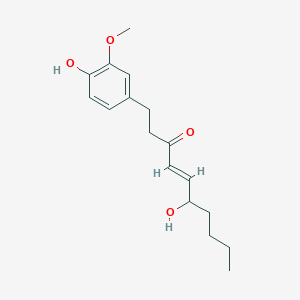

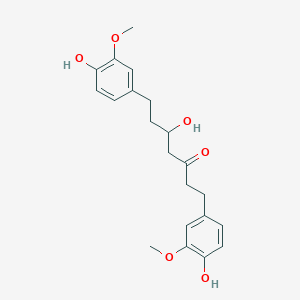

[6] – hydroxyshogaol	Hexahydrocurcumin

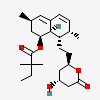

Simvastatin

**Figure S1** The chemical structures of selected ginger rhizome extracts and reference drug that retrieved from Pub-Chem Data bank

**Table S3** ADMET properties of screened ginger extract compounds computed using AdmetSAR webserver

| Molecule | 6-gingerol | 6- hydroxyshogaol | 8-shogaol | 10-gingerol | 10-shogaol |
| --- | --- | --- | --- | --- | --- |
| Log S solubility (mg/ml) | -3.23 | -2.85 | -5.80 | -6.42 | -6.92 |
| GI absorption | 0.9805 | High | High | High | High |
| BBB permeant | 0.6072 (yes) | Yes | yes | yes | Yes |
| Pgp substrate | 0.7319 (No) | No | No | yes | No |
| CYP1A2 inhibitor | yes | Yes | yes | yes | Yes |
| CYP2C19 inhibitor | No | No | No | No | No |
| CYP2C9 inhibitor | No | No | No | No | No |
| CYP2D6 inhibitor | yes | Yes | Yes | Yes | Yes |
| CYP3A4 inhibitor | No | No | Yes | Yes | Yes |
| Log Kp (cm/s)  (skin permeation) | -6.14 | -6.30 | -4.55 | -4.70 | -3.95 |

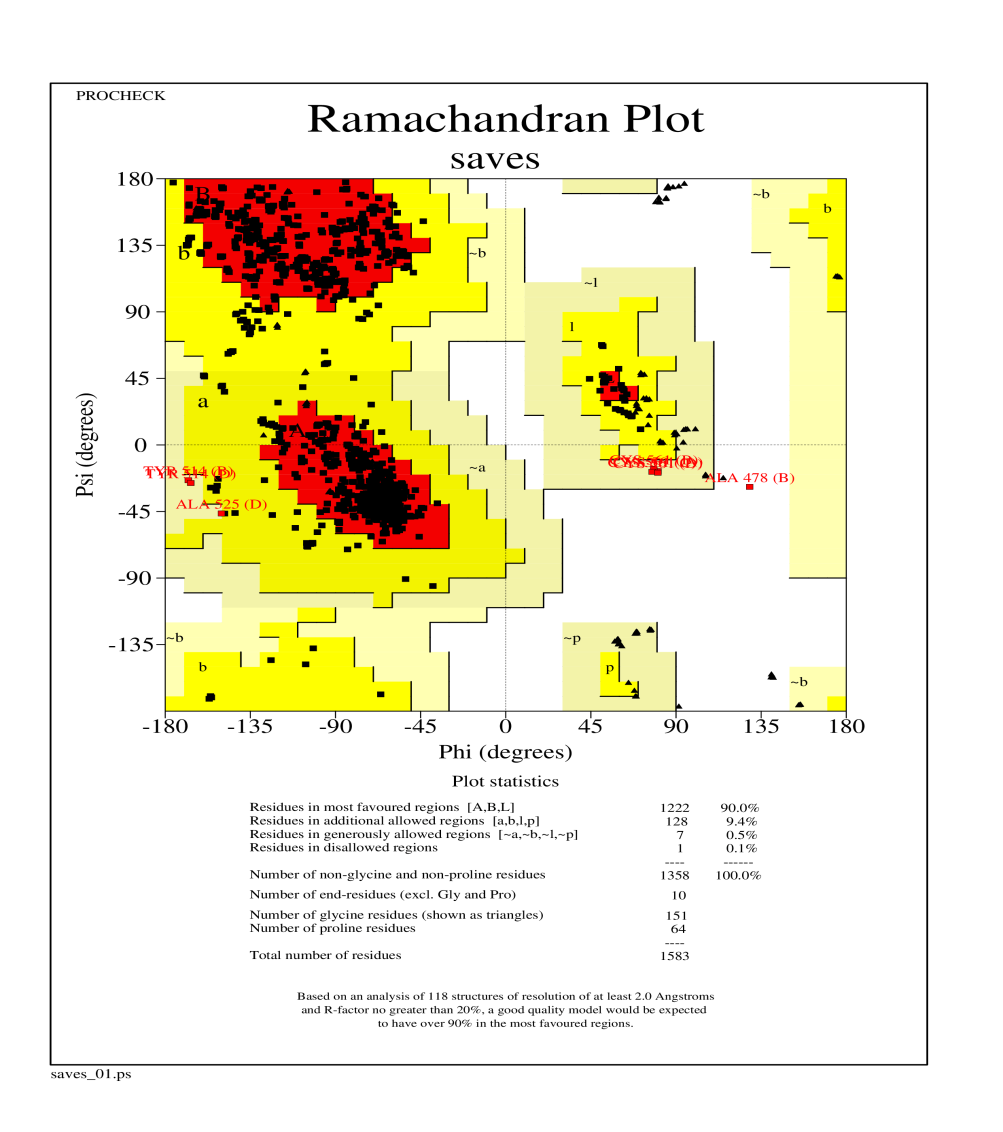

**Figure S2** Ramachandran plot showing the stereo chemical quality of the predicted HMG-CoA reductase (PDB ID: 1hw9) structure by using PROCHECK-DOE-MBI Structure Lab UCLA

**Reference**

1. Colovos C and Yeates TO. (1993). Verification of protein structures: patterns of nonbonded atomic interactions. *Protein Science*; 2(9):1511-9.
2. Pontius J, Richelle J and Wodak SJ. (1996). Deviations from standard atomic volumes as a quality measure for protein crystal structures. *J Mol Biol*; 264(1):121-36.
3. Laskowski RA, MacArthur MW, Moss DS and Thornton JM. (1993). PROCHECK a program to check the stereochemical quality of protein structures. *J. App. Cryst*; 26: 283-291.
4. Dong J, Wang NN, Yao ZJ, Zhang L, Cheng Y, Ouyang D, Lu AP and Cao DS. (2018). ADMETlab: a platform for systematic ADMET evaluation based on a comprehensively collected ADMET database. *Journal of Cheminformatics*; 10:29
